# Switching circuits: Modulating NK cell response to TGF-β using novel receptors

**DOI:** 10.1101/2023.12.10.570687

**Authors:** Rebekah Turk, Lan Guo, Brian J. Belmont, Nick Bogard, Aye T. Chen, Maddie D. Williams, Mary Young, Matthew R. Stone, Bryce Daines, Max Darnell

**Affiliations:** Modulus Therapeutics, Seattle, WA, USA

**Keywords:** TGF-β, chimeric switch receptor, B cell lymphoma, chimeric antigen receptor-T cell, chimeric antigen receptor NK cell

## Abstract

Engineered natural killer (NK) cells offer a promising treatment strategy for multiple therapeutic areas, yet tuning cell function to diverse environmental and inhibitory contexts remains a significant challenge. An attractive strategy for functional rescue in the presence of inhibitory ligands is the use of switch receptors, surface-expressed chimeric receptors that convert an external inhibitory cue into an intracellular activation signal. Here, we discover novel engineered switch receptors in NK cells that are responsive to TGF-β, a soluble inhibitory factor present in many therapeutic contexts. Through a pooled screen of an 11,131 member library, we identified multiple novel signaling endodomains and endodomain combinations that improve both the acute cytotoxicity and persistence of NK cells in the presence of TGF-β, demonstrating a novel and flexible approach to switch receptor discovery.

## INTRODUCTION

Engineered natural killer (NK) cells are increasingly being explored for their therapeutic potential as an alternative to T cell therapies. NK cells are more naturally amenable than T cells to allogeneic delivery without additional cell engineering, and have demonstrated a lower risk of serious side effects including cytokine release syndrome (CRS).^1–3^

NK cells are attractive templates for cell engineering because of their native anti-infective and anti-tumor function. NK cells produce immunostimulatory cytokines and chemokines, and are able to lyse cancerous or infected cells without prior sensitization.^1^ The initiation of these functions is dependent on the balance of activating and inhibitory receptors^2,3^, natural cytotoxicity receptors, and co-stimulatory receptors.^4^

Despite these advantages, NK cells, like T cells, are inhibited by a number of immunosuppressive mechanisms employed by target cells, including impairment of antigen presentation, activation of inhibitory costimulatory signaling pathways, and secretion of immunosuppressive factors.^5^ One potent example is hijacking Transforming Growth Factor-beta (TGF-β) regulation of immune responses by many types of tumors and in the context of certain viruses.^6,7^ TGF-β is a regulatory cytokine that plays an essential role in regulating differentiation, proliferation, migration and survival of immune cells and maintaining peripheral tolerance. Secretion of abundant TGF-β is a potent inhibitor of lymphocytes including NK cells, and attenuates cytotoxic activity and production of proinflammatory cytokines such as IFN-ɣ, an antagonist of IL-12 production in NKs.^8–11^

Overcoming inhibitory signals remains a significant challenge for cell therapies. Various engineering approaches have proliferated in recent years including an abundance of Chimeric Antigen Receptor (CAR) designs such as tandem CAR, dual-signaling CARs, AND-gate CARs, inhibitory CAR, AND-NOT CARs, CARs with three scFvs, ON/OFF-switch CARs, and universal CARs.^12^

Switch receptors are an emerging approach to engineering more functionality into NK cell therapies. Switch receptors combine the ectodomain of a cell-surface receptor to bind an inhibitory factor (e.g. TGF-βR2/TGF-β), and an activating endodomain (e.g. 4-1BB) to propagate proliferation and activating signals. A functional TGF-β switch receptor could transform inhibitory signals into activating signals enhancing NK cell proliferation and acute cytotoxicity in the presence of pathological levels of TGF-β. In contrast to dominant-negative receptors that bind to an inhibitory ligand but do not drive any signaling, switch receptors have the advantage of coupling beneficial signaling to the engagement of an inhibitory ligand. However, methods to characterize switch receptor designs in high throughput have been lacking, potentially limiting their therapeutic utility.

To expand the switch receptor design-space, we compiled a list of 105 endodomains from literature and functional motif-mining and designed a combinatorial library with 11,131 switch receptor constructs, each construct containing one or two endodomains. We utilized the high-throughput pooled screening methodology described previously^13^ and modified to select for TGF-βR2 switch receptors that increase proliferation and acute cytotoxicity in the presence of pathological levels of TGF-β. Our high-throughput screening strategy was designed to identify endodomains that drive enhanced persistence while maintaining high levels of cytotoxicity in an inhibitory context. This approach enables testing a large library of potential endodomain variants, and accelerates discovery by increasing the throughput of the experimentation. Improved understanding of the endodomains that influence NK persistence and cytotoxicity in the presence of TGF-β could lead to the engineering of NK cell therapies that are more effective within the context of, for example, solid tumors. Furthermore, beyond TGF-β, 6*this discovery method demonstrates the breadth of functionality and signaling diversity that can be driven by switch receptors, and motivates their expanded use in cell therapies.

## RESULTS

### Construction of a 11,131 member switch receptor library

The endodomain library consisted of 105 endodomains. We identified intracellular domains or functional motifs from immune receptors and signaling molecules (“native”, N=60) and made *de novo* endodomains by repeating select functional motifs two times (“synthetic”, N=44), one negative control sequence (“spacer”) was also included. This endodomain library expanded on previous endodomain libraries^13^ to include endodomains from receptors that activate NK cells along JAK/STAT-independent pathways and cytokine signaling pathways. The library also included domains from proteins involved in synapse formation, kinases, and cytoplasmic/intracellular signaling molecules.

Nested GoldenGate assembly was used to combinatorially generate an 11,131 member library of switch receptors. The TGF-βR2 switch receptor comprises the extracellular domain of TGF-βR2, the transmembrane domain of TGF-βR2 (or ROR2 in some control constructs), and one to two variable intracellular domains (endodomains) from a broad set of signaling molecules (Figure 1A). A lentivirus pool was made from the plasmid libraries and transduced into PBMC-derived NK cells.

**Figure 1.**
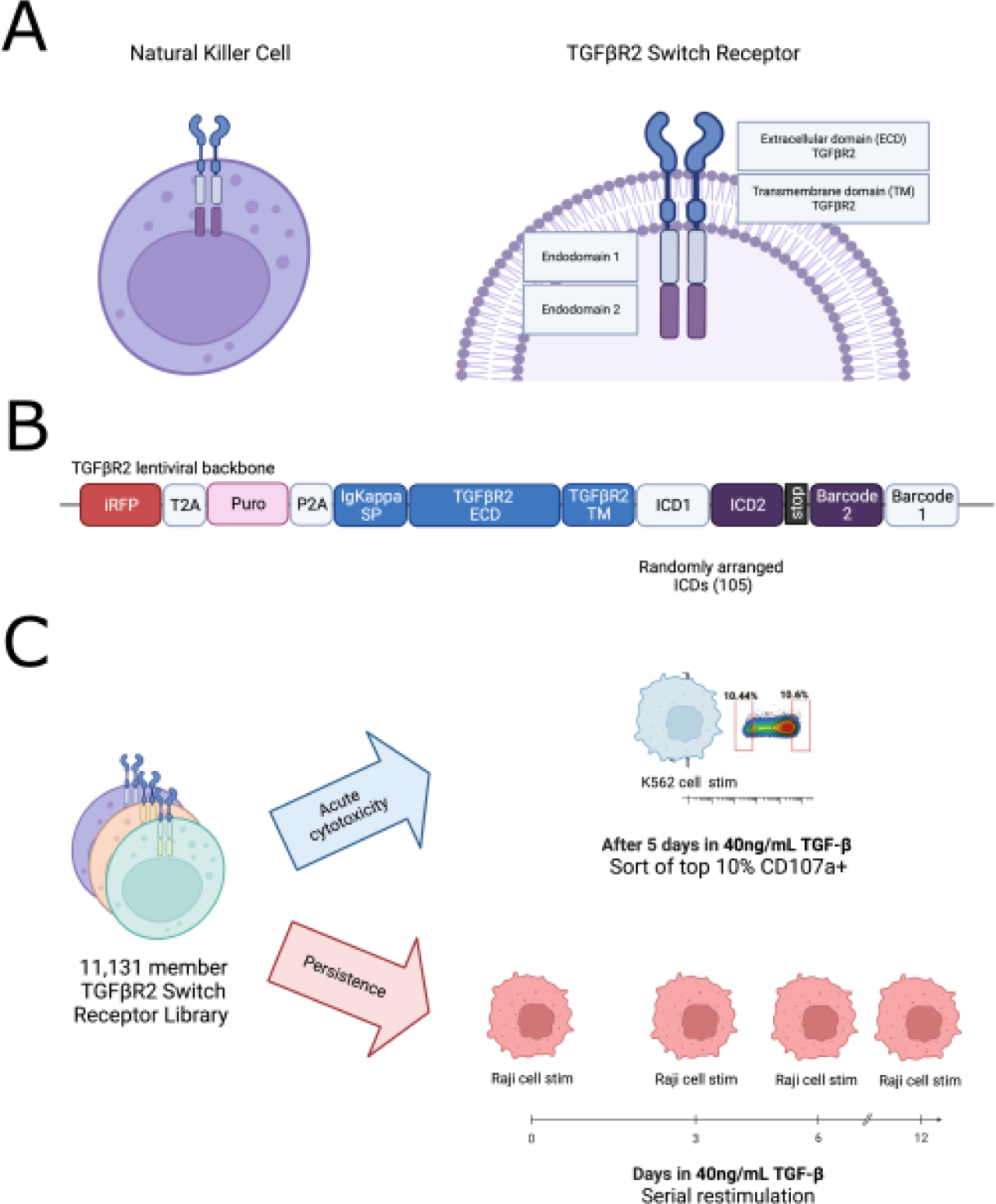
Switch receptor natural killer cells and TGF-βR2 switch receptor components. A) Illustration of TGF-βR2 switch receptor on NK cell surface, TGF-βR2 switch receptors consist of extracellular domain (ECD) that binds TGF-β, transmembrane domain (TM), and two intracellular domains or endodomains. B) Schematic of the lentiviral cassette used to express the switch receptor constructs. C) Schematic of the switch receptor library pooled screening workflow. Top: Acute cytotoxicity arm, bottom: Persistence arm.

### Switch receptors rescue TGF-β-induced suppression of NK persistence and cytotoxicity

It was previously demonstrated that exposing activated, expanded NK cells to pathologic levels of TGF-β leads to impaired function. Following a two-week expansion, NK cells maintained in culture with 5ng/ml or 10ng/ml TGF-β resulted in statistically significant impairment of cytotoxicity at 24h, 60h, and 96h post exposure compared to cells maintained without supplemented TGF-β.^14^ To optimize conditions for pooled screening workflows, select switch receptors were tested in varying concentrations of TGF-β (20 ng/mL or 40 ng/mL) added in the media for five days, with and without target cell co-culture. The following switch receptor constructs were included: TGF-βR2 ectodomain with ROR2 transmembrane domain and TGF-βR2 endodomain (TGFBR2_ROR2_TGFBR2); TGF-βR2 ectodomain with TGF-βR2 transmembrane domain and no endodomain (TGFBR2_TGFBR2_GG1); TGF-βR2 ectodomain with ROR2 transmembrane domain and endodomains from Fn14 and FCER1G (TGFBR2_ROR2_Fn14FCy); TGF-βR2 ectodomain with ROR2 transmembrane domain and endodomain from 4-1BB (TGFBR2_ROR2_4-1BB). TGFBR2_ROR2_TGFBR2 mimics the structure of the native TGF-β receptor whereas TGFBR2_TGFBR2_GG1 is designed to act as a TGF-β dominant negative as it only contains the ectodomain without any endodomain. Negative controls included a backbone-only construct and NK cells with no viral transduction (mock). Cell numbers and cell proliferation were quantified at 24 hours, 96 hours, and 120 hours post TGF-β exposure. Expression of NK phenotypic markers NKG2D, CD16, and CD107a was assessed through flow cytometry at each time point.

We observed that 40 ng/mL TGF-β inhibits the growth of NK cells compared to no TGF-β in the media from 3-5 days in culture (Supplementary Figure 1A). This effect was more pronounced when the receptor was overexpressed (TGFBR2_ROR2_TGFBR2, red line) and in the mock condition (purple line), which had lower ratio of cell counts with TGF-β to cell counts without TGF-β. The TGFBR2_ROR2_Fn14FCy construct had the highest ratio of cell counts with TGF-β to cell counts without TGF-β at day 5, which suggested endodomain signaling through the switch receptor could be counteracting the TGF-β suppression of growth. TGF-β treatment also decreased the cytotoxicity of NK cells in a dose-dependent manner, as evidenced by the increased numbers of live Raji cells co-cultured with NK cells in 20 ng/mL and 40 ng/mL TGF-β compared to no TGF-β (Supplementary Figure 1B). Compared to the dominant negative receptor TGFBR2_TGFBR2_GG1, both TGFBR2_TGFBR2_4-1BB (magenta line) and TGFBR2_TGFBR2_Fn14FCy (orange line) achieved better killing (lower live Raji counts) in the presence of TGF-β (Supplementary Figure 1C), with TGFBR2_TGFBR2_Fn14FCy out-performing TGFBR2_TGFBR2_4-1BB. This is consistent with the expression of the cytolytic marker CD107a after 5 days in culture, assessed with flow cytometry (Supplementary Figure 1D), as both TGFBR2_TGFBR2_4-1BB (magenta dot) and TGFBR2_TGFBR2_Fn14FCy (orange dot) showed a higher percentage of CD107a-positive cells than the dominant negative receptor TGFBR2_TGFBR2_GG1.

The largest effects on cell proliferation and phenotype marker expression were observed with 4-5 days of 40 ng/mL TGF-β exposure. Based on these data, we selected the dose of 40 ng/mL TGF-β for 5 days as the protocol, and CD107a as the cytotoxic marker to sort on for the pooled screening.

### Switch receptor endodomain pooled screens reveal novel functional constructs and endodomain combinations

Following the optimization work, we proceeded to screen the full switch receptor library in a pooled setting. Aiming to discover constructs that promote both cytotoxicity and persistence in suppressive TGF-β conditions with antigen challenge, our workflow employed two distinct arms - an ‘acute cytotoxicity’ arm and a ‘persistence’ arm. For the acute cytotoxicity arm, the switch receptor NK cell library was cultured in media with and without 40 ng/mL TGF-β for 5 days, and then stimulated with K562 target cells for 3 hours. The cells were then stained with anti-CD107a and sorted by flow cytometry to collect the top 10% and bottom 10% CD107a expressing cells. For the persistence arm, NK cells were cultured in media with and without 40 ng/mL TGF-β and mitomycin-C-treated target Raji cells were added at an E:T of 1:1 every 3 days for 12 days to provide repeated NK activation cues. Amplicon sequencing was used in both workflows to quantify switch receptor enrichment. DESeq2 enrichment analysis was performed on construct count data to compare construct abundance with and without TGF-β treatment in the acute cytotoxicity workflow and before and after prolonged antigen stimulation with TGF-β treatment in the persistence workflow.

TGF-β exposure and antigen stimulation posed a strong selective pressure on the switch receptor pool. In the acute cytotoxicity arm, comparing construct counts in the top 10% CD107a-positive NK cells treated with 40 ng/mL TGF-β for 5 days to the top 10% CD107a-positive NK cells without TGF-β yielded 491 significantly enriched (log2 fold change > 0 and adjusted p < 0.05) and 404 significantly depleted (log2 fold change < 0 and adjusted p < 0.05) constructs (Figure 2A). In the persistence arm, comparing NK cells after 12 days of antigen rechallenge with 40 ng/mL TGF-β exposure to those without TGF-β exposure yielded 1687 significantly enriched and 1447 significantly depleted constructs (Figure 2B).

**Figure 2.**
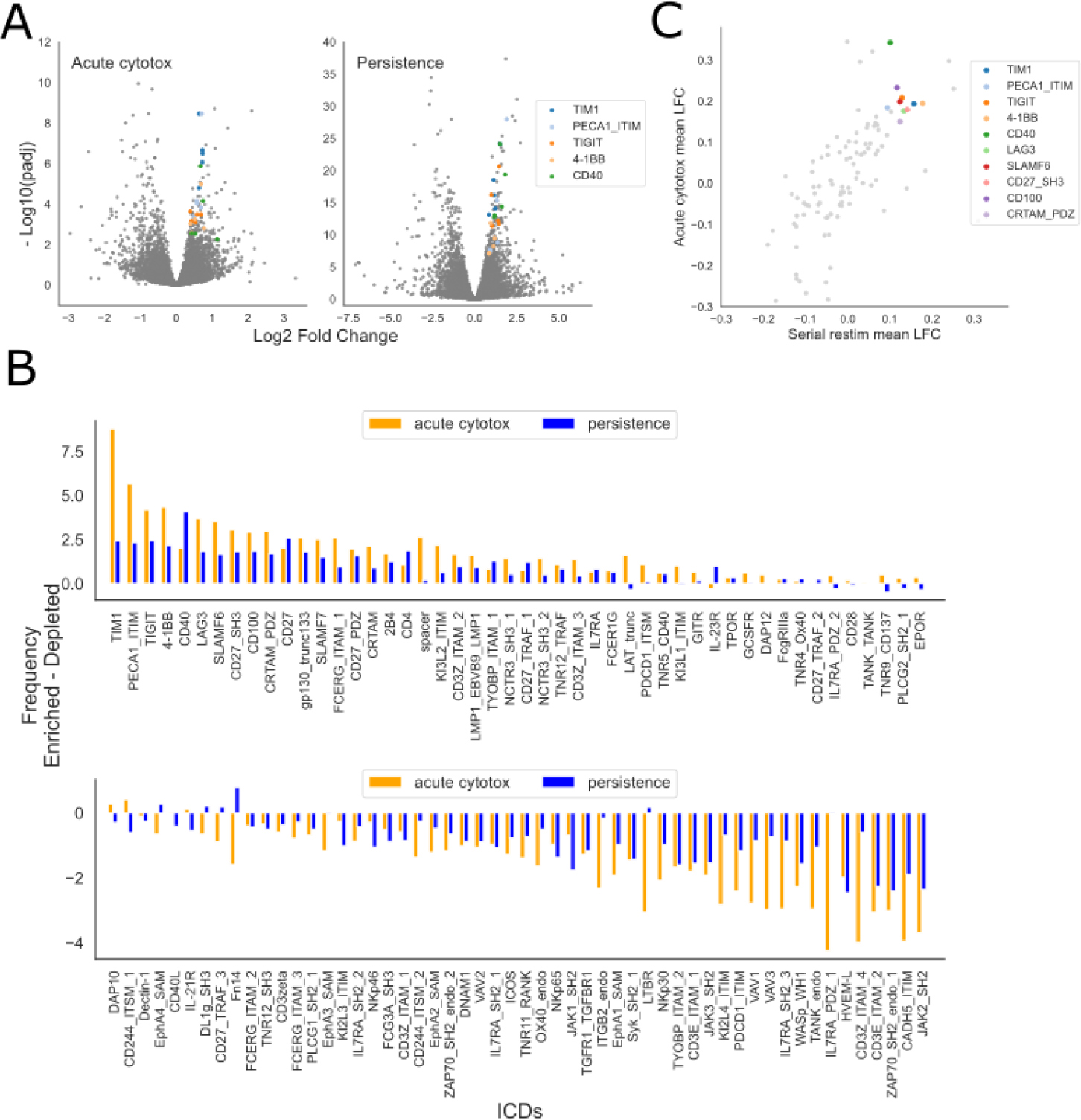
TGF-βR2 switch receptor library pooled screen results. A) Volcano plots of DESeq2 results for acute cytotoxicity (left) and persistence (right) workflows of the pooled switch receptor screen, highlighting top enriched endodomains (colored dots). DESeq2 was used to calculate the log2 fold changes and BH-adjusted p-values for each construct, comparing NK cells sorted by CD107a expression (top 10% CD107a-positive cells) treated with 40 ng/mL TGF-β vs. no TGF-β. Endodomains that were present in high frequency in the top enriched constructs were colored. B) Bar plot for each individual endodomain, showing its frequency in significantly enriched constructs minus its frequency in significantly depleted constructs in both acute cytotoxicity workflow (orange) and persistence workflow (blue). Top: endodomains that are over-represented in significantly enriched constructs; bottom: endodomains that are over-represented in significantly depleted constructs. C) Mean LFC for all constructs containing each individual endodomain in acute cytotoxicity workflow vs. persistence workflow, highlighting top endodomains (colored dots).

To determine which signaling components convey an advantage against the selective pressure in both workflows, we investigated which endodomain components were observed at higher frequencies in significantly enriched constructs following both the acute cytotoxicity and persistence protocols (Figure 2C). Since these endodomains were combinatorially assembled into pairs during library generation, each can occur at either position and in combination with all the other endodomain library members. We calculated the average fold change for all constructs that contain an individual endodomain and found these to be well correlated (Pearson’s R=0.65, p<0.001) between the acute cytotoxicity workflow and the persistence workflow (Figure 2D). A subset of endodomains (colored dots) are enriched for both acute cytotoxicity and persistence workflows, these include both the canonical co-stimulatory 4-1BB domain but also novel endodomains such as CD40, PECA1_ITIM, gp130_trunc133, TIGIT, and TIM1.

### Novel switch receptors validated for persistence and killing in arrayed format

To validate the observed effects observed in our pooled screen, we performed arrayed validation experiments using 40 candidates that spanned a range of enrichment values in the pooled screen. A backbone-only construct was included as negative control; a construct with 4-1BB endodomain was included as positive control. Lentiviruses expressing each construct were separately prepared to transduce NK cells, and all constructs were expanded following enrichment for switch receptor expressing cells. These candidates were assessed for their cytotoxic function against K562 target cells after growing in media supplemented with 40 ng/mL TGF-β for five days.

The cytotoxicity index for each switch receptor candidate was calculated (see Methods for details). Switch receptors that were enriched (log2FoldChange > 0) in the pooled screen acute cytotoxicity arm demonstrated a higher cytotoxicity index compared to 4-1BB and switch receptors that were depleted (log2FoldChange < 0) in the pooled screen acute cytotoxicity arm (Figure 3A, Wilcoxon rank sums test p = 0.001). Specifically, ranking the switch receptors by the mean cytotoxicity index of the replicates, 15 of the top 20 vs. 6 of the bottom 20 switch receptors were enriched in the pooled screens. Ten switch receptors from our pooled screen exhibited better performance (higher cytotoxicity index) than the positive control 4-1BB. In addition, NK cell growth was quantified after 5 days of culture in media with and without 40 ng/mL TGF-β. Pooled screen enriched switch receptors performed slightly better at counteracting the TGF-β-induced inhibition of growth than pooled screen depleted switches (Wilcoxon rank sums test p > 0.05), although all switch receptors tested appear to convey growth enhancing effect in the presence of TGF-β over that of the backbone-only control (Figure 3B).

**Figure 3.**
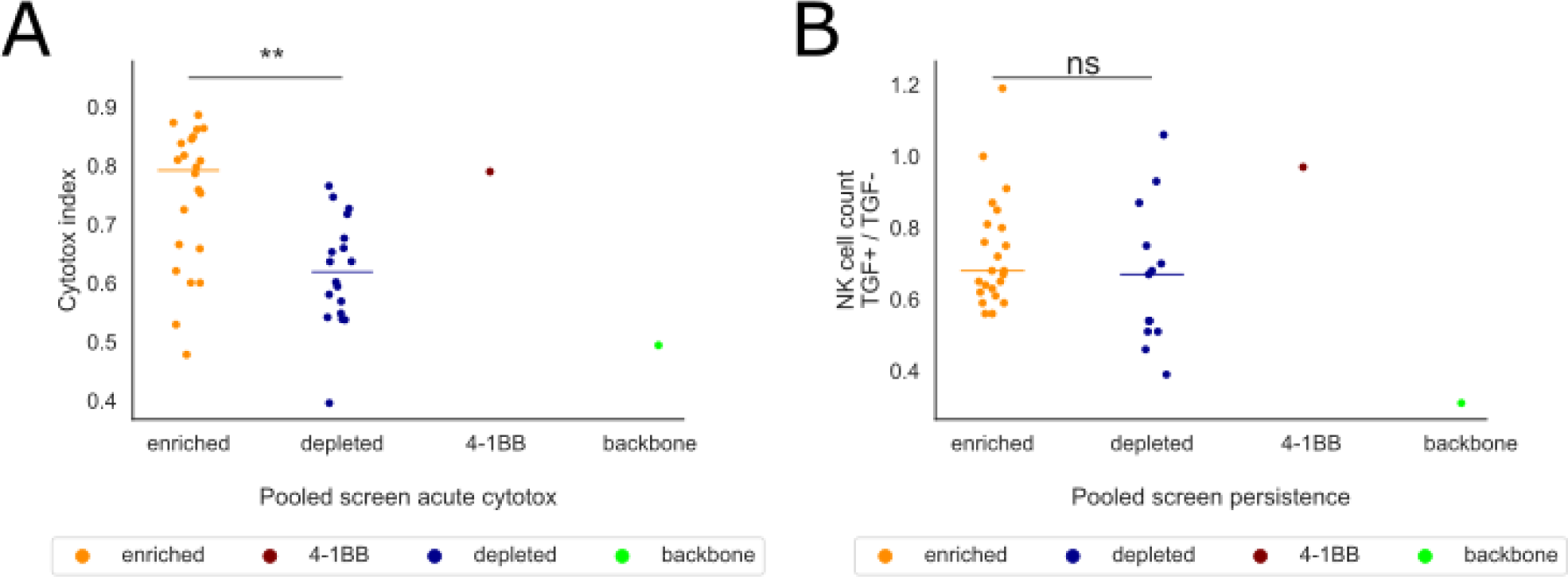
Arrayed validation of TGF-βR2 switch receptor designs spanning a range of pooled screen performance. A) Forty two switch receptor constructs were validated for acute cytotoxicity in the arrayed format by growing in 40 ng/mL TGF-β-containing media for five days followed by incubating for three hours at an E:T ratio of 2:1 against 4e4 K562 target cells. The cytotoxicity index was calculated as (1 - # target cells remaining in NK-containing wells / # target cells remaining in target-only well). Dot plot shows average cytotoxicity index from 2 technical replicates for each construct. Colors indicate constructs that were enriched (log2 fold change > 0), depleted (log2 fold change < 0), the positive control 4-1BB, and the backbone-only negative control. Horizontal colored lines indicate median values for enriched and depleted constructs. **: Wilcoxon rank sums test p < 0.001. B) Live NK cell counts were quantified via flow cytometry at the end of five days in culture with media containing 40 ng/mL TGF-β or no TGF-β. Dot plot shows the cell count of the 40 ng/mL TGF-β condition normalized to that of no TGF-β condition for each construct. Colors indicate constructs that were enriched (log2 fold change > 0), depleted (log2 fold change < 0), the positive control 4-1BB, and the backbone-only negative control. Horizontal colored lines indicate median values for enriched and depleted constructs. ns: Wilcoxon rank sums test p > 0.05.

### Top hit selection via arrayed acute cytotoxicity and spheroid assays

Based on the results from the initial validation effort, we sought to assess a larger set of switch candidates that were enriched in the pooled screen in order to identify highly performing sequences. For this set of experiments, we selected 78 positively enriched constructs from the acute cytotoxicity arm and 74 positively enriched constructs from the persistence arm of the pooled screen, along with 4-1BB as the positive control and a backbone-only construct as the negative control. Experiments were conducted in an arrayed format similar to the initial validation. These candidates were assessed for their acute cytotoxic function against K562 target cells after growing in media supplemented with 40 ng/mL TGF-β for five days. Cytotox index was calculated as described in the initial validation experiments. Forty-four out of 152 (28 out of 74 selected from the persistence arm and 16 out of 78 selected from the acute cytotoxicity arm) of our positive hits performed better than 4-1BB in lysing K562 target cells after TGF-β exposure (Figure 4A).

**Figure 4.**
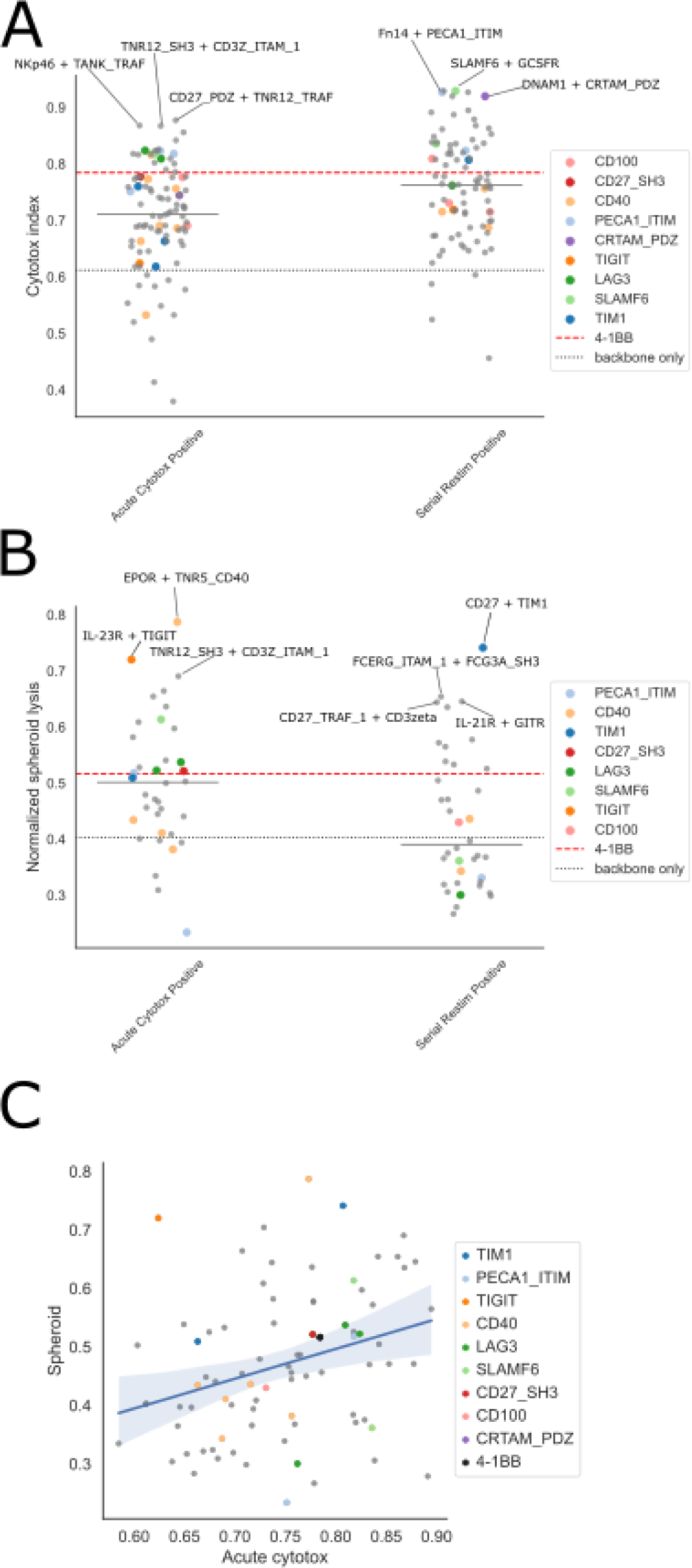
Acute cytotoxicity and persistent killing of spheroid cultures. A) 78 positively enriched constructs from the acute cytotoxicity arm and 74 positively enriched constructs from the persistence arm of the pooled screen were validated for acute cytotoxicity in the arrayed format by growing in 40 ng/mL TGF-β-containing media for five days followed by incubating for four hours at an E:T ratio of 2:1 against 1e4 K562 target cells. The cytotoxicity index was calculated as (1 - # target cells remaining in NK-containing wells / # target cells remaining in target-only well). Dot plot shows average cytotoxicity index from 2 technical replicates for each construct. Red dashed line indicates the average performance of the positive control 4-1BB, and black dotted line indicates the backbone-only negative control. Horizontal gray lines indicate median values for each category. Color dots represent constructs that contain a specified endodomain. B) 74 positive hits (38 selected from the persistence arm and 36 selected from the acute cytotoxicity arm) were tested for chronic cytotoxicity by incubating with GFP-expressing Raji cells at an E:T ratio of 3:1 for 100 hours on a BioTek Cytation 5 with BioSpa 8 and imaged every 2 hours. The area under the curve (AUC) for the GFP integral signal over the whole duration of the experiment for each well was calculated as the total spheroid growth. The normalized spheroid lysis was calculated as (1 - AUC in NK-containing well / mean AUC in target-only well). Dot plot shows average cytotoxicity index from 2 technical replicates for each construct. Red dashed line indicates the average performance of the positive control 4-1BB, and black dotted line indicates the backbone-only negative control. Horizontal gray lines indicate median values for each category. Color dots represent constructs that contain a specified endodomain. C) Correlation of cytotox index and normalized spheroid lysis in 79 switch constructs tested in both A) and B). Color dots represent constructs that contain a specified endodomain.

To increase the stringency of the screening process, we additionally performed time-lapse imaging in a Raji spheroid model to assess the prolonged cytotoxic potential of the candidates. NK cells expressing switch constructs were incubated with GFP-expressing Raji cells at an E:T ratio of 3:1 for 100 hours on a BioTek Cytation 5 with BioSpa 8 and imaged every 2 hours. The average GFP integral signal over time from the detected spheroids was extracted. To estimate the total spheroid growth overtime, we calculated the area under the curve (AUC) for the GFP integral signal over the whole duration of the experiment for each well. The AUCs for the target-only wells on each plate were averaged and used for normalization to calculate the spheroid lysis for wells containing NKs expressing the switch constructs (see Methods). Out of the 74 positive hits tested (38 selected from the persistence arm and 36 selected from the acute cytotoxicity arm), 27 (11 from the persistence arm and 16 from the acute cytotoxicity arm) outperformed 4-1BB (Figure 4B).

Although the spheroid assay yielded more variable results than the acute cytotoxicity assay, the two assays’ results were generally consistent. For the switch constructs that were tested in both assays (N=79), their acute and chronic cytotoxic measurements were positively correlated (Figure 4C, Spearman’s R=0.34, p=0.0019).

## DISCUSSION

We introduce a novel method for high throughput switch receptor discovery that offers a flexible strategy for precisely tuning cell therapy function with a single receptor architecture. As an example, we use this method to demonstrate an approach to engineer CAR NK cells to overcome a key functional bottleneck to effective solid tumor clearance: the inhibitory cytokine, TGF-β. Our screening scheme identified promising switch receptor candidates that improve acute cytotoxicity and persistence of CAR NK cells in the presence of TGF-β beyond industry benchmarks. Several of our top hits feature novel domains not presently used in cell therapies. Our results suggest that this discovery approach can be used to identify switch receptor structural components tailored to specific cells signaling cues.

We sought to discover switch receptors to enhance not only NK acute cytotoxicity but also persistence. Despite the associated screens being run in parallel and focused on distinct functions, we observed a high degree of correlation between the results for individual switches within the screen (Fig. 2C). The connection between the cellular dynamics that underlie cytotoxicity and persistence are not well understood, but our results may suggest some degree of overlapping signaling pathways. Furthermore, the variation that is observed between the switches along both functional axes provides a multitude of options for the selection of a switch receptor well-suited to any particular cell design or clinical indication, allowing the cell engineer to ‘program’ the cell from a library of functions.

The molecular mechanism of switch receptor activation was not evaluated here, but the validated switches included domains that are capable of two distinct modes of signal transduction: 1) mechanotransduction (e.g. 4-1BB, CD40^15^) or 2) homodimerization (e.g. IL-7R^16^). Canonically, CARs and switch receptors have relied on mechanotransduction-based mechanisms to initiate signal transduction. This mechanism generally relies upon the interaction of a surface-bound or immobilized ligand with a cell-surface receptor to impart a mechanical force on the receptor, leading to receptor signal activation. In contrast, dimerization-based receptor signal transduction relies upon ligand-receptor interactions leading to the formation of heterodimer or homodimer complexes, which trigger the signaling cascade.

We hypothesize that it is the dimeric form of TGF-β that enables this soluble ligand to engage with both, unrelated forms of signaling architecture. For the mechanotransduction case, TGF-β has previously been shown to trigger activation in T cells by binding two separate switch receptors in *cis* on the cell surface, leading to the requisite force transmission on one another^17^. The binding of two separate switch receptors in *cis* could also colocalize the receptors, leading to homodimer-based signaling. One exemplary library member is the IL-7R endodomain, which has been shown to activate in homodimer complexes^16^ and demonstrates activity in the context of switch receptors described here. A TGF-β/IL7R switch receptor has been previously reported in T-cells, although the precise signaling mechanisms were not thoroughly evaluated^18^. Collectively, these models of signal transduction predict a lack of switch receptor activity in the presence of a monomeric soluble factor, and therefore the endodomain architectures of the switch receptors discovered here may not generalize to other soluble ligands.

The screening approach described here has the potential to be employed broadly across cell types, therapeutic contexts, and switch receptor ligands. While switch receptors canonically have relied on inhibitory receptors as the basis for receptor design, the functional space of switch receptors is large. For example, in a scenario in which NK cells are already activated against a target cell, switch receptors targeting non-inhibitory receptors could enhance cell fitness, promote cell-cell contact, and/or stabilize immune synapse formation leading to a spectrum of positive effects. Furthermore, switch receptors are a flexible engineering tool that have the potential to be deployed in complementary sets in order to tune cell function. Engaging with the multi-parametric design space of a cell featuring multiple switch receptors, with each receptor featuring multiple signaling domains, will require high throughput discovery approaches such as those described here.

## MATERIALS & METHODS

### Design of switch receptor library

We compiled a list of 105 endodomains which expanded on libraries previously reported in CAR endodomain screening^13^. The expanded endodomains include 60 derived from endogenous proteins, 44 designed from duplicating signaling motifs, and one negative control consisting of a non-signaling spacer motif. Grouping endodomains based on their downstream binding partners and pathways, there were 22 TRAF binding, 34 Syk/ZAP-70 associated, 20 PI3K/Fyn/Lck binding, 15 JAK-STAT binding, 5 synapse formation, and 9 inhibitory or miscellaneous proteins. Since we discovered TANK as a hit in our previous CAR endodomain study, we included a few other cytoplasmic/intracellular signaling molecules such as VAV1.

### Assembly of library by GoldenGate

Switch receptor library was assembled as described previously^13^. We used nested GoldenGate assembly to generate a combinatorial library with 11,131 switch receptor constructs each containing one to two endodomains. We ordered two sets of 105 endodomains and their specific barcodes as eBlocks from IDT. In GoldenGate step 1, we used the first set of eBlocks to clone a pool of 105 single endodomain constructs. In GoldenGate step 2, we cloned the second set of 105 eBlocks into the pool of constructs from step 1. Since GoldenGate cloning had residual background, the final library contained 11,025 (105 x 105) double endodomain constructs, 105 single endodomain constructs, and one original backbone construct (11,131 total unique constructs).

### Cell lines

Cell lines were maintained as described previously^13^ with the addition of Gibco Viral Production 2.0 cells (Thermo Fisher). These were cultured in Gibco Viral Production Media supplemented with 4mM GlutaMAX (Thermo Fisher) and kept shaking at 125 RPM on a 19 mm orbital shaker at 8% CO2.

### Lentivirus generation (screen)

Lentivirus for the pooled screens was generated as described previously^13^.

### High Throughput Lentivirus generation (validation only)

Gibco Viral Production Cells 2.0 HEK293 suspension cells were grown in Gibco Viral Production Media and passaged in a Thompson Ultra Yield Flask to maintain confluency under 6e6 cells/mL. The day before transfection, VPC 2.0 cells were seeded at a density of 2.1e6 cells per mL.

For transfection, cells were aliquoted at a volume of 3 mL/well into 24-well pyramid-bottomed plates. A DNA mass of 2 ug DNA/mL of cells was used at a molar ratio of 3:1:1:1 for the transfer plasmid, GagPol RRE, Rev, and pMD2.G plasmids, respectively. Transfections were carried out using Xfect (Takara) following the manufacturer’s instructions and using an Xfect to DNA ratio of 0.3 uL:1 ug per mL of cells.

Three days post-transfection, viral supernatant was harvested by pelleting cells at 1500xg for 10 minutes. The viral particles in the supernatant were then concentrated using LentiX Concentrator (Takara Bio), in accordance with the manufacturer’s instructions and resuspended at 10X final concentration in PBS.

### Primary NK cell isolation

Human NK cells were isolated directly from a fresh quarter-leukopak by CD56-negative immunomagnetic enrichment (STEMCELL #17955, EasySep™ Human NK Cell Isolation Kit) and were frozen down in CS10 CryoStor cell cryopreservation media media (Sigma-Aldrich).

### Primary NK cell expansion

NK cells were cultured and expanded as described previously^13^.

### Primary NK cell transduction

Feeder-cell expanded NK cells were transduced in duplicate 10 days post-thaw. Thirty minutes prior to transduction, NK cells were pretreated with 3 uM BX-795 (Sellek). Pretreated NK cells, lentivirus, polybrene (Sigma-Aldrich) at a final concentration of 8 ug/mL and BX-795 at a final concentration of 3 uM were combined and spinoculated in a 96-well U-bottom plate in a centrifuge at 1200xg for 30 minutes at 32°C. After spinoculation, the cells were resuspended and incubated at 37°C overnight. Post incubation, cells were media exchanged and transferred to a GREX plate of appropriate size (Wilson Wolf, New Brighton Minnesota, USA).

### Switch NK cell enrichment

Three days post-transduction, the TGF-β switch NK cells were fed with mitomycin C-treated K562 feeder cells at a 1:1 ratio of feeder cells:NK cells. Four days post-transduction, puromycin selection was performed by adding puromycin (InvivoGen) at a final concentration of 2 ug/mL. The TGF-β switch NK cells were maintained in puromycin for 5 days until fully selected and then media exchanged into puromycin-free media.

### Flow cytometry

Transduction efficiency was estimated using iRFP670 contained within the switch plasmid. Raji cells were estimated by LSSmOrange marker. Human Fc receptors were blocked using Human TruStain FcX (Biolegend). Phenotyping was performed on a NovoCyte Advanteon flow cytometer and FACS was performed on a BD LSRII.

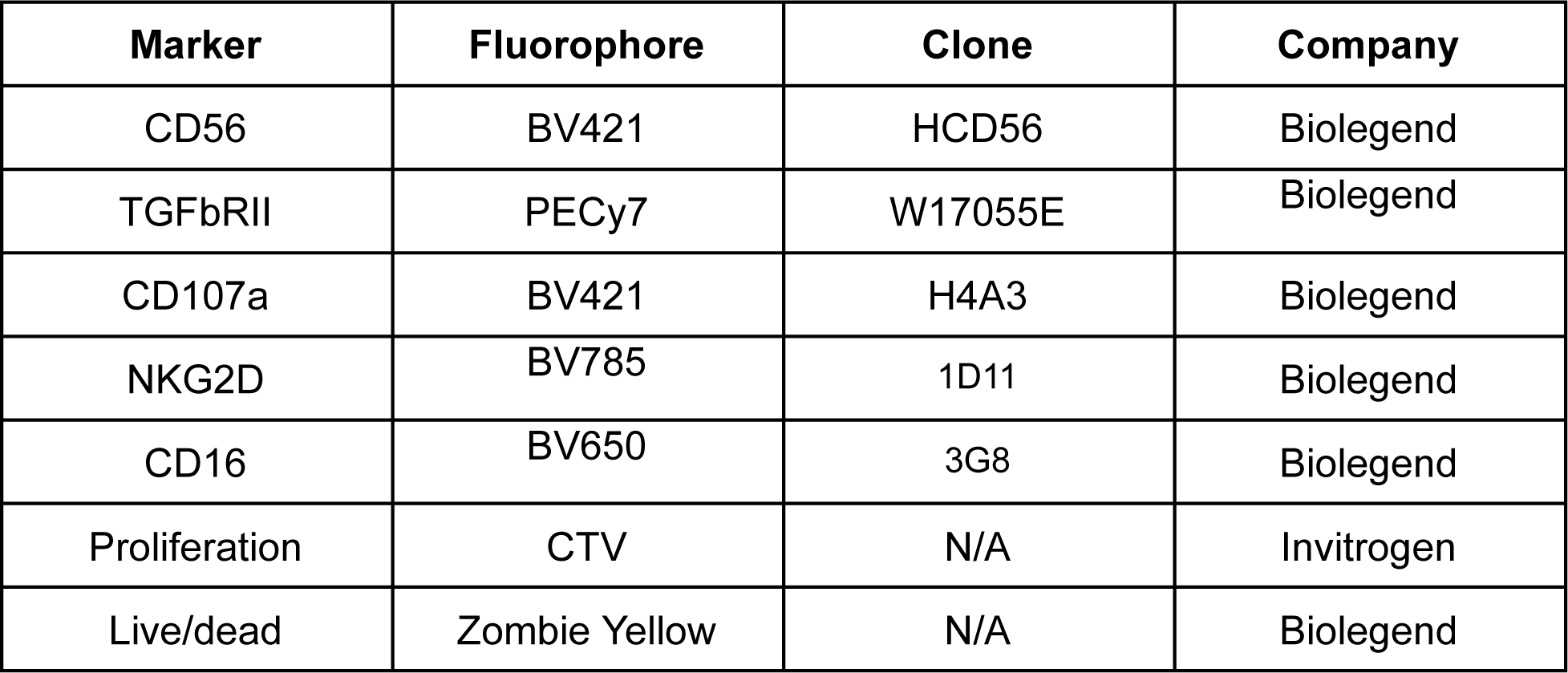

### Serial restimulation screen

For the pooled co-culturing assay, the TGF-β switch NK cell library experienced four different assay conditions for 12 days: 1) no TGF-β without cells, 2) no TGF-β with cells, 3) plus TGF-β without cells, and 4) plus TGF-β with cells. Generally, six days after feeding, post-puromycin selected TGF-β switch library transduction replicates were plated at 5e6 cells/well in a six-well GREX plate. For the conditions with cells, mitomycin-C treated (to inhibit growth) Raji cells were added to achieve a 1:1 ratio of TGF-β switch library cells:Raji cells every three days, starting at day 0. For conditions plus TGF-β, TGF-β switch library cells were grown in media supplemented with or without 40 ng/mL TGF-β. Approximately 1e6 cells per condition, in duplicate, were collected directly into RLT buffer supplemented with 38 mM DTT every three days for genomic DNA (gDNA) isolation, with amplicon sequencing performed on the day 0 and day 12 collection points.

### Acute cytotoxicity screen

For the pooled acute cytotoxicity assay, the TGF-β switch NK cell library was sorted via FACS using CD107a expression profiles after target cell coculture. Six days after feeding, post-puromycin selected TGF-β switch NK cell library transduction replicates were cultured in media supplemented with or without 40 ng/mL TGF-β for 5 days. TGF-β switch NK cells were then cocultured with K562 GFP cells at an E:T ratio of 1:1 in the presence of 1X monensin and anti-CD107a antibody for three hours and sorted on CD107a high and CD107a low, as defined by the top and bottom 10% of each population, respectively. Approximately 1e6 cells per condition, in duplicate, were collected directly into RLT buffer supplemented with 38 mM DTT followed by amplicon sequencing.

### Acute cytotoxicity screen validation

Forty candidates that spanned a range of predicted activities were generated in the same manner as the TGF-β switch NK cell library and incubated in media supplemented with 40 ng/mL TGF-β for five days. These cells were then co-cultured in a 96-well round-bottomed plate for four hours at an E:T ratio of 2:1 against 4e4 K562 GFP target cells and the cytotoxicity index was calculated (1 - # target cells remaining after 3 hours coculture/# target cells remaining in target-only well), with cell counts determined via a NovoCyte Advanteon flow cytometer.

### Top hits validation

Putative top hits selected from both the acute and serial restimulation workflows were generated in the same manner as the TGF-β switch NK cell library and cultured in media supplemented with and without 40 ng/mL TGF-β for five days. These cells were then sent through two assay conditions: acute cytotoxicity and spheroid killing. The acute cytotoxicity assay replicated the acute cytotoxicity screen validation, with an E:T ratio of 2:1, but with a reduced target cell number of 1e4 K562 cells. For the spheroid assay, Raji GFP were cells seeded in 96-well ultra-low attachment plates (S-BIO PrimeSurface® 3D Culture Spheroid plates #MS-9096UZ) at 1e4 cells/well 24 hours prior to coculture in media supplemented with or without 40 ng/mL TGF-β. NK cells were then added on top of the spheroids at an E:T of 3:1. The plates were incubated at 37°C and 5% CO2 in a BioTek BioSpa 8, with the average GFP integral captured via a BioTek Cytation 5 on 465nM LED excitation, SYBR Green emission every 2 hours for 100 hours. The average GFP integral signal from all spheroids detected in a well at each timepoint was exported. To determine total spheroid growth overtime, the area under the GFP integral time course was calculated using the trapz function from the numpy package. An average of the target-only wells was determined as the mean NK-free spheroid growth. For all wells containing NK cells, the normalized spheroid lysis was calculated as (1 - AUC in NK-containing well / mean AUC in target-only well).

### Illumina sequencing

Cells were lysed in RLT buffer (QIAGEN) supplemented with 38 mM DTT. Genomic DNA was isolated and Illumina sequencing was subsequently performed as described previously^13^. We sequenced both the endodomain and barcode sequences within the initial plasmid library to determine the barcode swapping rate between barcode1 and barcode2 during GoldenGate cloning. We observed less than 4% barcode swapping between the two barcodes (more than 96% of the Illumina sequencing reads contained correct endodomain and barcode pairs). Samples from the screen were sequenced for barcodes only. Sequencing was done on either Illumina iSeq or NextSeq 2000 Systems.

### Processing and analysis of NGS data

Processing and analysis of NGS data was performed as described previously^13^.

### HT DNA Preparation

To prepare DNA for the arrayed validation, 10 uL of bacterial glycerol stocks was innoculated into 1 mL of TB media containing 100 ug/mL carbenicillin in 96-well 2 mL deep-well blocks, and cultured overnight in a shaking incubator at 900 rpm and 37°C. Overnight cultures were harvested and DNA purified using QIAGEN QIAprep 96 Turbo Kits following the manufacturer’s instructions. DNA concentrations were quantified using UV plates and the Agilent BioTek Cytation 5 Multimode Reader.

## SUPPLEMENTARY FIGURES

**Supplementary Figure 1.**
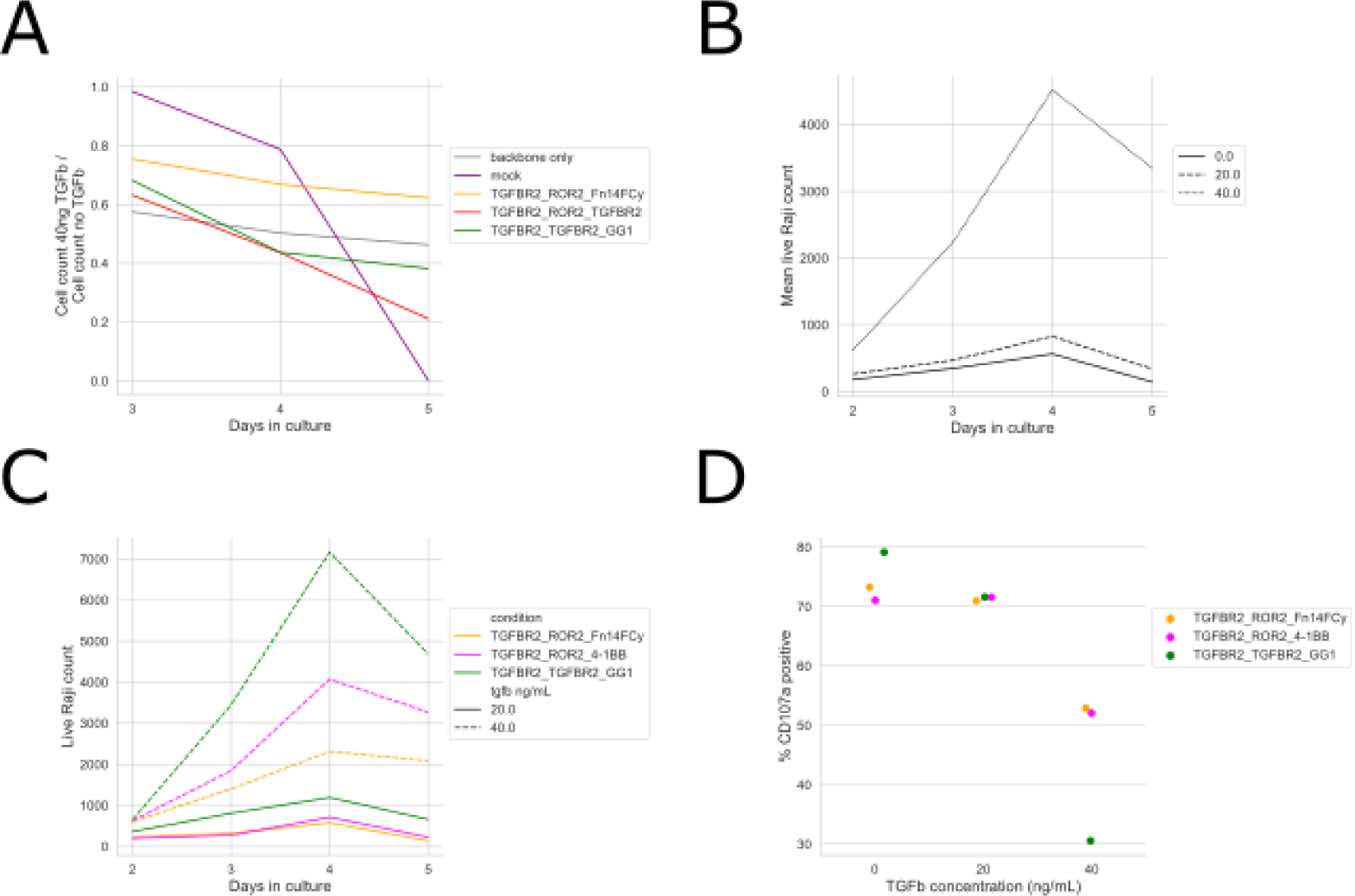
TGF-βR2 switch receptors rescue TGF-β-induced suppression of NK persistence and cytotoxicity. A) NK cell count in 40 ng/mL TGF-β divided by NK cell count in no TGF-β from 3-5 days in culture. B) Mean counts of live Raji cells co-cultured with NK cells in 20 ng/mL and 40 ng/mL TGF-β compared to no TGF-β for all constructs. C) Live Raji cell counts for TGFBR2_TGFBR2_GG1 (green), TGFBR2_TGFBR2_4-1BB (magenta) and TGFBR2_TGFBR2_Fn14FCy (orange) in 20 ng/mL and 40 ng/mL TGF-β. D) The percent of CD107a-positive cells for each construct after 5 days in culture with 0, 20, and 40 ng/mL TGF-β, assessed with flow cytometry.

**Table 1.**
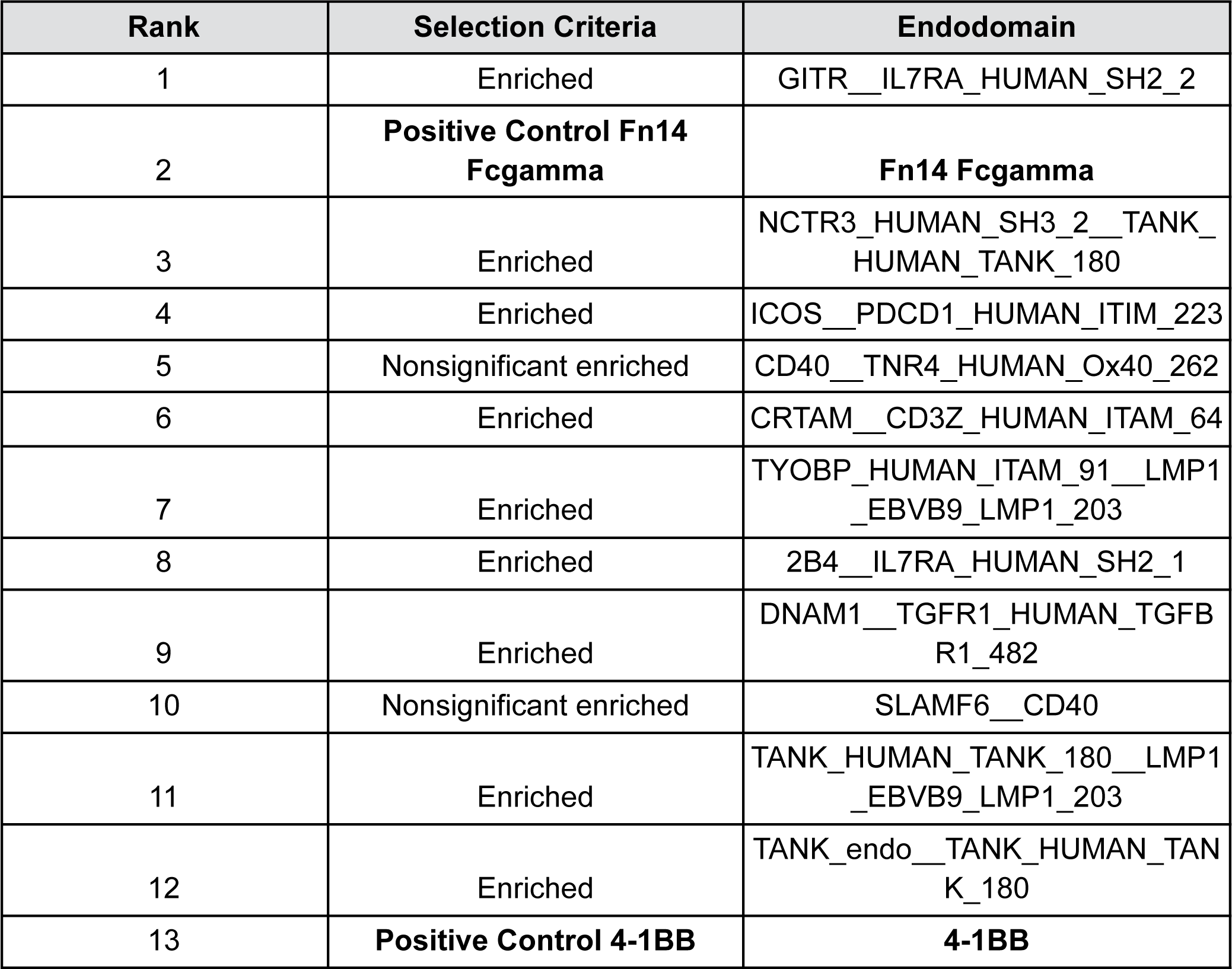
Endodomains validated by colony picking with cytotoxicity indexes higher than 4-1BB.

